# Visualization of viral infection dynamics in a unicellular eukaryote and quantification of viral production using VirusFISH

**DOI:** 10.1101/849455

**Authors:** Yaiza M. Castillo, Marta Sebastián, Irene Forn, Nigel Grimsley, Sheree Yau, Cristina Moraru, Dolors Vaqué

## Abstract

One of the major challenges in viral ecology is to assess the impact of viruses in controlling the abundance of specific hosts in the environment. For this, techniques that enable the detection and quantification of virus–host interactions at the single-cell level are essential. With this goal in mind, we implemented VirusFISH (Virus Fluorescence *in situ* Hybridization) using as a model the marine picoeukaryote *Ostreococcus tauri* and its virus OtV5. VirusFISH allowed the visualization and quantification of the fraction of infected cells during an infection experiment. We were also able to quantify the abundance of free viruses released during cell lysis and assess the burst size of our non-axenic culture, because we could discriminate OtV5 from phages. Our results showed that although the major lysis of the culture occurred between 24 and 48 h after OtV5 inoculation, some new viruses were produced between 8 and 24 h, propagating the infection. Nevertheless, the production of viral particles increased drastically after 24 h. The burst size for the *O. tauri–*OtV5 system was 7±0.4 OtV5 per cell, which was consistent with the estimated amount of viruses inside the cell prior to cell lysis. With this work we demonstrate that VirusFISH is a promising technique to study specific virus–host interactions in non-axenic cultures, and set the ground for its application in complex natural communities.

## INTRODUCTION

Marine viruses have been studied during the last three decades, mostly by traditional approaches as microscopy (Noble and Fuhrman, 1998) and flow cytometry (Marie *et al.*, 1999), used for the enumeration and estimation of viral production. However, in the last few years, the development of high throughput sequencing techniques has considerably changed the field, and our knowledge about viral communities has exponentially increased. These new sequencing approaches provide information about the viral taxonomic and genomic diversity, about their biogeography and, to a certain extent, about their potential hosts (*e.g.* Chow *et al.*, 2015; Labonté *et al.*, 2015). However, they do not allow the visualization of specific virus–host interactions and the monitoring of the infection dynamics, which are crucial to better understand the contribution of viruses in shaping microbial communities and biogeochemical cycles. Attempts to identify virus–host associations date back to the 90s, when the role of viruses in the marine environment started to be recognized. Hennes *et al.* (1995) pioneered an approach to identify and enumerate specific virus-infected bacteria in natural communities by using fluorescently stained viruses (labeled with YOYO-1 or POPO-1) as probes and epifluorescence microscopy. Years after, Tadmor *et al.* (2011) used microfluidic digital PCR to detect specific phage–host associations in the termite gut. With this method, they managed to directly detect the phage–host association by targeting genes from both components without culturing, but with no visual representation of the infection.

A few years ago, Allers *et al.* (2013) developed phageFISH and used it to monitor phage infections at the single-cell level in a marine podovirus—gammaproteobacterial host system. PhageFISH uses mixtures of polynucleotide probes labeled with digoxigenin to target phage genes, and a single HRP labeled oligonucleotide probe to target host rRNA. The signal from the two types of probes is amplified and visualized by catalyzed reporter deposition (CARD) of fluorescently labeled tyramides. Compared to the method from Hennes *et al.*, (1995), where the infection was forced by adding stained viruses to identify the host within natural communities, phageFISH enables the visualization of the infection dynamics of specific virus—host pairs, because it simultaneously targets the virus and the host. More recently developed, direct-geneFISH (Barrero-Canosa *et al.*, 2017) uses simultaneously a mixture of polynucleotide probes directly labeled with fluorochromes, to detect specific genes in cells, and a single oligonucleotide probe, carrying multiple fluorochromes, to identify bacterial cells.

In the present work, we combined the phageFISH and direct-geneFISH techniques to develop VirusFISH with the aim to allow i) identification and quantification of specific virus—unicellular eukaryote interactions at the single-cell level and ii) identification and quantification of free virus particles. VirusFISH consists of two steps. First, a CARD-FISH step is used to detect host cells, with HRP-labeled oligonucleotide probes targeting the 18S rRNA. Then, a VirusFISH step is applied to detect viruses, using multiple polynucleotide probes directly labeled with fluorochromes that target viral genes. VirusFISH can be used to detect both intracellular viruses and free viral particles.

As proof of principle, we used VirusFISH to monitor viral infections of the unicellular green alga *Ostreococcus tauri* (*O. tauri*), the smallest known marine photosynthetic eukaryote, with the virus *Ostreococcus tauri* virus 5 (OtV5).

## MATERIALS AND METHODS

### Experimental viral infection of *O. tauri*

The host strain *O. tauri* RCC4221 (Roscoff Culture Collection, NCBI accession number txid70448) was grown in 60 mL of L1 medium (Guillard and Hargraves, 1993) in aerated flasks (Sarstedt), and incubated at 21.5°C (±0.5°C) with white light ∼100 µE and a 10:14 hours photoperiod (light:darkness), until stationary phase (7.16×10^7^ ± 3.57×10^6^ cells mL^-1^, estimated by 4′-6-Diamidino-2-phenylindole (DAPI) counts). Triplicate *O. tauri* cultures (20 mL) were infected with 1 mL of OtV5 inoculum (1.3×10^7^ ± 4.3×10^6^ viruses mL^-1^, estimated by plaque-forming units), resulting in a 0.01 MOI (multiplicity of infection). Non-infected triplicate *O. tauri* cultures (inoculated with 1 mL of L1 medium) were used as control. After OtV5 inoculation, samples (900 µL) were taken over 3 days at times 0, 8, 24, 48 and 72 h, and fixed with 100 µL of freshly filtered formaldehyde (3.7% final concentration) for 15 min at room temperature. Then, 500 µL of fixed sample were filtered through 0.2 µm pore size polycarbonate white filters (Merck™ GTTP02500) to retain cells, and through 0.02 µm pore size anodisc filters (Whatman®) (after a 0.2 µm pore size prefiltration to remove cells and debris) to retain free viruses. Polycarbonate filters of 0.2 µm pore size were embedded in 0.1% (w v^-1^) low gelling point agarose and treated for 1h with 96% ethanol and 1h with pure methanol, to remove cellular pigments that can interfere with the CARD-FISH signal (Fig. S1), and 10 min with HCl to inactivate endogenous peroxidases (Pavlekovic *et al.*, 2009). All filters were kept at -20°C until hybridization.

### OtV5 probe design and synthesis

For the detection of the OtV5 virus (NCBI accession number EU304328) we designed 11 dsDNA polynucleotide probes (300 bp each) using the software geneProber web service (http://gene-prober.icbm.de/). These 11 probes covered a total of 3998 bp of the OtV5 viral genome, offering sufficient sensitivity to detect single viruses (Table S1), as it has been shown before (Barrero-Canosa *et al.*, 2017). Each probe synthesis was done by obtaining the corresponding polynucleotides by PCR, and then all probes were mixed and labeled with the Alexa594 fluorochrome, based on the protocol from Barrero-Canosa *et al.* (2017). The PCR was set up as follows: 10pg of OtV5 DNA were added to a reaction mixture containing 200 µM (each) deoxyribonucleoside triphosphates (Invitrogen), 1 µM of each primer, 1x PCR buffer (Invitrogen), and 5U of *Taq* DNA polymerase (Invitrogen). The thermal cycling was performed in a C1000TM Thermal Cycler (Bio-Rad) with an initial denaturation step at 95°C (5 min), followed by 30 rounds at 95°C (1 min), X°C (30 s), and 72°C (30 s), and a final extension at 72°C (10 min). X value corresponds to the optimal annealing temperature for each of the primers, determined after performing gradient PCRs. All OtV5 primers had an optimal annealing temperature of 62.5°C, with the exception of primers #3 and #5 that had an annealing temperature of 65.5°C. Primers sequences can be found in Table S1. For each polynucleotide, several PCRs were done to obtain a minimum of 400µL PCR reaction volume. This volume was purified on a single purification column using the QIAquick PCR purification kit – Qiagen, cat.no. 28106, and resuspended in a TE solution (5 mM Tris-HCl, 1 mM EDTA, pH 8.0). The polynucleotide length was checked by agarose gel electrophoresis, and the concentration was measured spectrophotometrically using a NanoDrop 1000 (Fisher Thermo Scientific). Further, all 11 polynucleotides were mixed equimolarly to yield a total of 1 µg DNA in 10 µl TE. Later, the probe mixture was heated to 95°C for 5 min to denature it and then incubated for 30 min at 80°C with 10 µL of the red emission dye Alexa594 (Ulysis™ Alexa Fluor® 594 Nucleic Acid Labeling Kit, Thermofisher, cat.no: U21654). The unbound Alexa594 was removed using chromatography columns (Micro Bio-spin chromatography columns P-30, Bio-Rad, cat.no. 732-6202). The concentration of the probe mixture and the labeling efficiency with Alexa 594 were determined spectrophotometrically using a NanoDrop 1000 with the Multi-Array option and N-50. For a successful detection of the virus, we observed that the labeling efficiency should be higher than 6 Alexas per probe. Fluorescent probes were stored at -20°C until use.

### Detection of *O. tauri* cells using 18S rRNA targeted CARD-FISH

*O. tauri* cells were labeled using Catalyzed Reporter Deposition-FISH (CARD)-FISH (Pernice *et al.*, 2015) with the 18S rRNA targeted probe OSTREO01 for *Ostreococcus* spp. (Not *et al.*, 2004). Briefly, the hybridization was carried out by covering filter pieces with 20 μL of hybridization buffer with 40% deionized formamide and incubating at 35 °C overnight. After two successive washing steps of 10 min at 37 °C in a washing buffer and a equilibration in phosphate-buffered saline for 15 min at room temperature (Cabello *et al.*, 2016), the signal was amplified for 1h at 46°C with Alexa488-labeled tyramide. Filters were then placed in phosphate-buffered saline two times for 10 min, rinsed with MilliQ water and air-dried.

### Detection of intracellular and free OtV5 viruses using VirusFISH

OtV5 viruses were labeled using VirusFISH, a modified version of the direct-geneFISH protocol (Barrero-Canosa *et al.*, 2017). VirusFISH was applied on i) 0.2 µm pore size filters that were previously hybridized with the CARD-FISH probes for the host to monitor the infection, ii) 0.02 µm pore size filters to monitor the dynamics of the free OtV5 viruses. The hybridization was done by covering the filter pieces with 25 µL of 40% formamide hybridization buffer (HB) containing OtV5 probes and incubating first for 40 minutes at 85°C, and then for 2 h at 46°C. The composition of the hybridization buffer was: 40% formamide, 5x saline-sodium citrate, 20% dextran sulfate, 0.1% sodium dodecyl sulfate, 20 mM EDTA, 0.25 mg mL^-1^ sheared salmon sperm, 0.25 mg mL^-1^ yeast RNA and 1% blocking reagent). The volume of probe mixture labeled with Alexa594 to add to the HB was calculated based on the following formula, according to Barrero-Canosa *et al.* (2017):

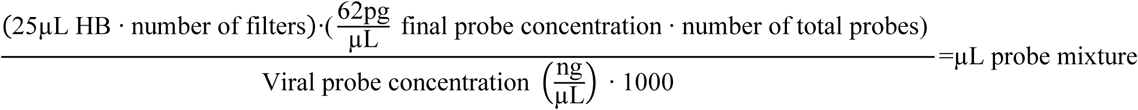

Assuming that the volume of HB for each filter portion is 25 µL and 62 pg µL^-1^ is the desired final probe concentration according to Barrero-Canosa et al. (2017).

Finally, samples were washed at 48°C for 15 minutes with gentle shaking in a washing buffer (560 µL NaCl 5M, 1mL Tris-HCl 1M pH8, 1mL EDTA 0.5M pH8 and 50µL 10% sodium dodecyl sulfate in 50mL of autoclaved MilliQ water), rinsed with MilliQ water and air-dried.

### Sample mounting, visualization and image analysis

After hybridization, 0.2 µm filters were counterstained with DAPI at 0.5 µg mL^-1^ to observe *O. tauri* nuclei, and mounted in antifading reagent (77% glycerol, 15% VECTASHIELD and 8% 20x PBS) (Cabello *et al.*, 2016). Images were manually acquired using a Zeiss Axio Imager Z2m epifluorescence microscope (Carl Zeiss, Germany) connected to a Zeiss camera (AxioCamHR, Carl Zeiss MicroImaging, S.L., Barcelona, Spain) at x1000 magnification through the AxioVision 4.8 software. *O. tauri* was observed by epifluorescence microscopy under blue light (475/30 nm excitation, 527/54 BP emission, and FT 495 beam splitter) and OtV5 under orange light (585/35 nm excitation, 615 LP emission, and FT 570 beam splitter). All pictures were taken using the same intensities and exposure times (400 ms for the *O. tauri* and 1 s for the virus detection).

Total free viruses (i.e. both OtV5 and phages present in the non-axenic culture) collected on the 0.02µm pore size filters, were counterstained with SYBRGold (SYBR™ Gold solution, Invitrogen) at 2x final concentration for 12 min, and then rinsed abundantly with MilliQ water to remove excess staining. Filters were finally mounted on slides with an antifading mounting solution (CitiFluor™ Glycerol-PBS Solution AF1). Images were acquired on the same Zeiss microscope and camera at x1000 magnification. OtV5 were observed by epifluorescence microscopy under orange light (585/35 nm excitation, 615 LP emission, and FT 570 beam splitter) and total viruses (OtV5 and phages) under blue light (475/30 nm excitation, 527/54 BP emission, and FT 495 beam splitter). All pictures were taken using the same intensities and exposure times as mentioned above. Image analysis for free virus detection was done using the software ACMEtool 3 (July 2014; M Zeder, Technobiology GmbH, Buchrain, Switzerland).

During the image analysis we observed that a fraction of OtV5 virions released from the cells during lysis was trapped on the extracellular organic matrix around the cells (here referred to as viral clouds) (Weinbauer *et al.*, 2009), and retained on the 0.2 µm filters. Thus, for 48 and 72 hours, we calculated the viral abundance of OtV5 retained on the 0.2 µm polycarbonate filters from the average area of viral clouds corrected by the average area of an OtV5 virus (48 h, n=2432 viral clouds areas; 72 h, n=307 viral clouds areas; OtV5, n=30,000 OtV5 particles areas). Consequently, we considered the total OtV5 production at 48 and 72 hours as the sum of the free virus abundance collected onto the 0.02 µm filters plus the viral abundance retained on the 0.2 µm. Areas were determined using the AxioVision 4.8 software (Schindelin *et al.*, 2012).

### Burst size estimations

Burst size was calculated based on the formula established by Middelboe and Lyck, (2002) that compares the Δvirus abundance / Δhost abundance at the times when the host decline happens (here, between 24 and 48 hours). To corroborate the burst size by the classical method, we also assessed the area of the host occupied with OtV5 at late stages of the infection. Since Henderson *et al.* (2007) reported that the structure of *Ostreococcus* is rather flattened, we calculated the average area of *O.tauri* hosting the OtV5 virions at 24 h (n=90 areas), when the maximum infection was observed, and the average area of single free OtV5 particles (n=30,000 viruses). The capacity was finally estimated by dividing the average cellular area occupied with viruses by the average area of a single viral particle. Areas were determined using the AxioVision 4.8 software (Schindelin *et al.*, 2012).

## RESULTS

### The OtV5 – *O. tauri* infection dynamics as revealed by VirusFISH

A non-axenic culture of *O. tauri* was infected with the virus OtV5, at a MOI of 0.01 and an uninfected culture was grown in parallel, as a control (Fig. 1A). Using VirusFISH, the two cultures were followed for 72 h, quantifying i) the absolute abundance of *O. tauri* cells and ii) the relative and absolute abundance of infected *O. tauri* cells. The infected culture experienced a dramatic decrease in cell density of two orders of magnitude between 24 and 48 h (Fig. 1B, Fig. 2 and Fig. S2). At 72h, almost no *O. tauri* cells were detected (Fig. S2), consistent with the clearing of the infected culture (Fig. 1A).

**Figure 1.**
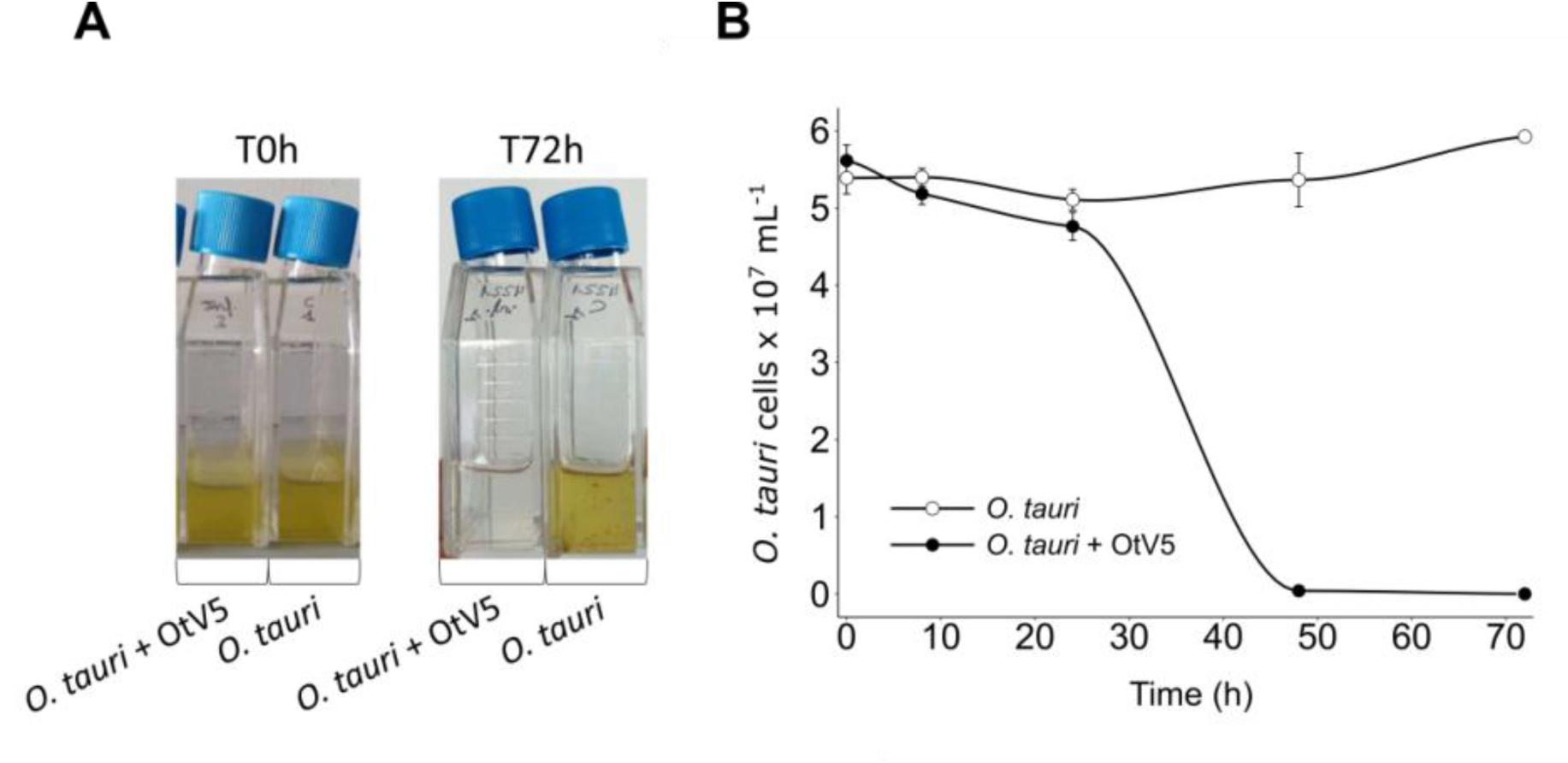
Dynamics of the infection of *Ostreococcus tauri* with OtV5. **A.** Infection and control culture flasks at time 0h and 72h. **B.** *O. tauri* CARD-FISH cell abundances (cells × 10^7^ ml^-1^) counted by epifluorescence microscopy in both the infected(solid circles) and the control (empty circles) triplicate cultures.

**Figure 2.**
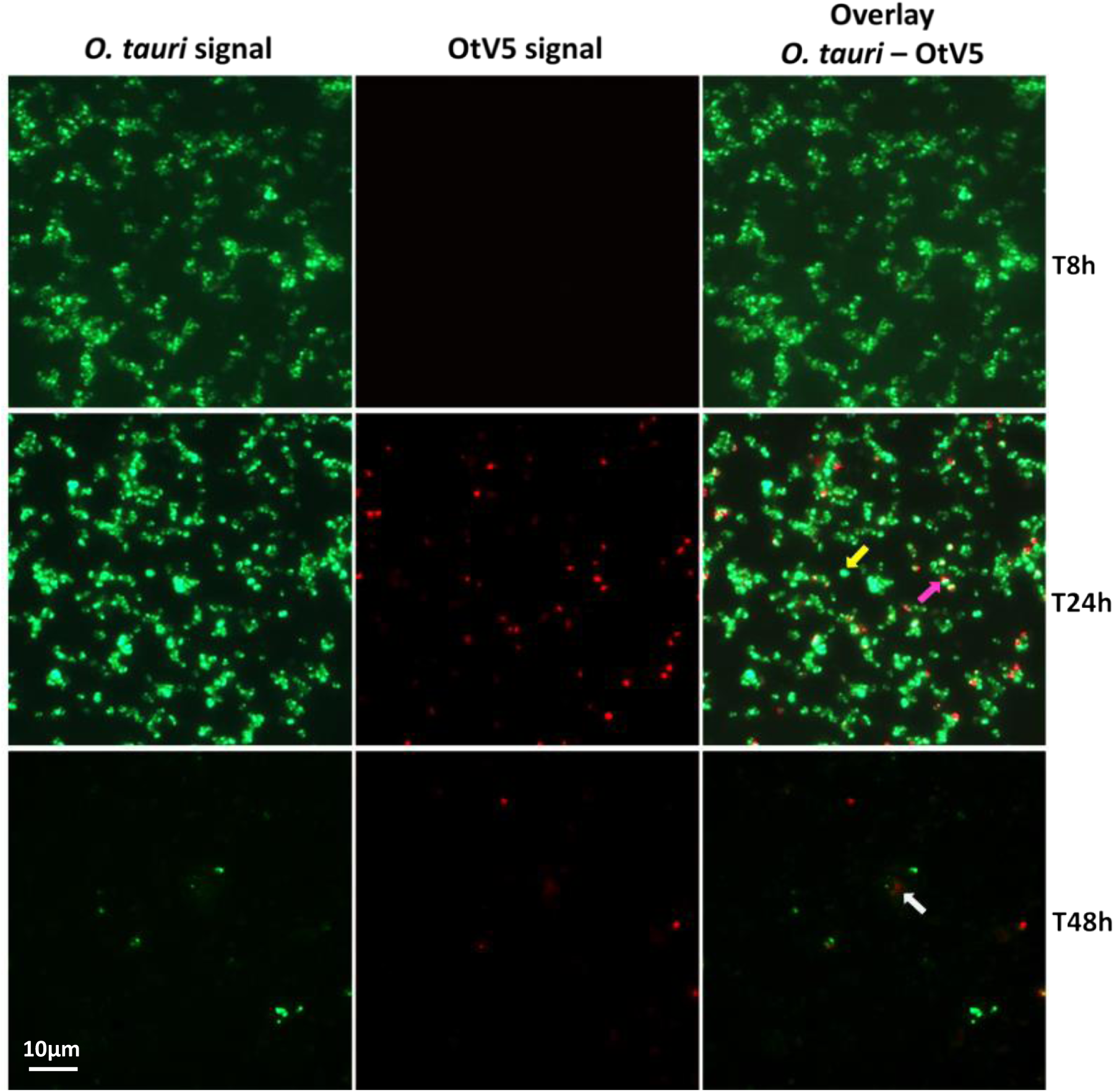
Micrographs of the evolution of the infection from time 8h to 48h. Left: *O. tauri* only. Centre: OtV5 only. Right column: overlay of *O. tauri* host cells in green (Alexa488) and virus in red (Alexa594). Yellow arrow: non-infected *O. tauri*; pink arrow: infected *O tauri*; grey arrow: cloud of viruses retained on the filter by the organic matter released during the lysis.

At the MOI used, rapid adsorption of all the viral particles added would theoretically result in 1% of infected cells. However, despite infected cells were visible as early as 0.4 h, the abundance was very low at both 0.4 and 8 h (0.02% and 0.2%, respectively), suggesting that not all viral particles had yet been adsorbed (Fig. 3). Nevertheless, the fact that at 24 h we found 16% of the population infected implies that new viruses had already been produced that had gone on to infect more cells in the culture. Later, at 48 h, the abundance of cells decreased by two orders of magnitude and 60% of the remaining cells were infected (Fig. 2 and 3). In contrast, the abundance of *O. tauri* cells in the control cultures remained relatively constant along the experiment and, as expected, no infected cells were observed (Fig. 1B and S3).

**Figure 3.**
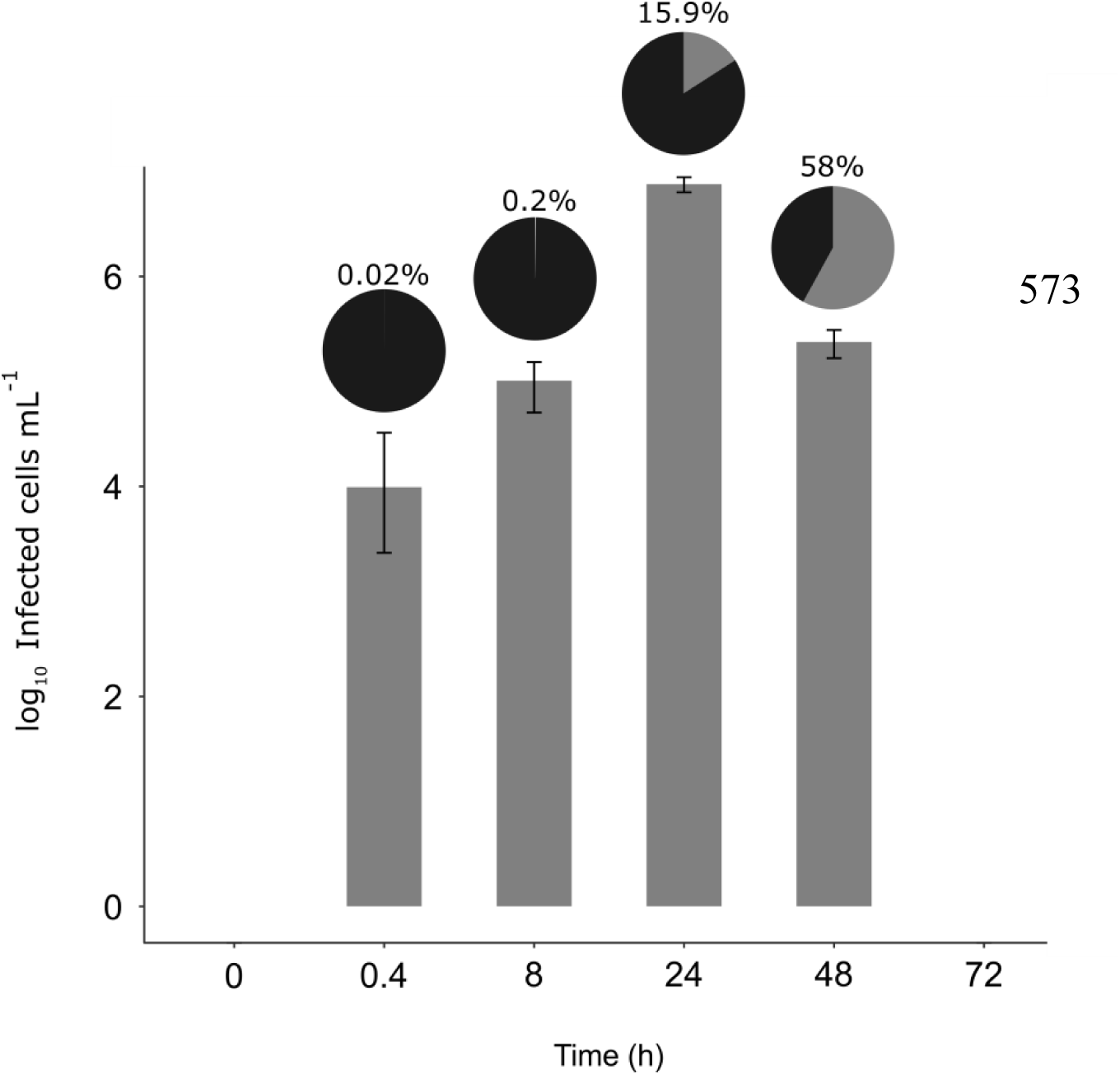
Dynamics of the infected cells. Bar plot shows the number of infected *O. tauri* at each time. Pie charts on top of each bar show the percentage of infected cells with respect to the total *O. tauri* abundance.

### Dynamics and abundances of free OtV5 particles

We also used VirusFISH for the detection and quantification of free OtV5 particles produced during the infection and lysis of *O. tauri*. As mentioned above, the *O. tauri* culture is not axenic, so we performed a SYBRGold staining step to label all the dsDNA viruses present (green particles in Figure 4), which include OtV5, bacteriophages, vesicles and/or other artefacts of non-specific staining. Since a certain background can be observed in the micrographs, only the VirusFISH red signal that overlapped with a SYBRGold green fluorescence signal was considered a true OtV5 particle (yellowish particles in Fig. 4). Our results showed that before 24 h, at least 1 infection cycle had already completed, as indicated by the slight, but detectable increase of OtV5 free particles and the slight decrease of *O. tauri* cells at 24 h. This is in agreement with the detection of 16% infected *O. tauri* cells at 24 h, which is higher than expected for the MOI used, as explained above. A drastic increase in viral abundance was observed after 24 h (Fig. 5 and Fig. S4), corresponding with the time the majority of cells were lysed. At 48 h the number of free viruses reached a plateau, likely because most viral production had already occurred. As expected, no increase in OtV5 particles was detected in the control flasks. The fraction of OtV5 within the total viral community labeled with SYBRGold ranged from 0.9% (±0.2%) at 0 h when viruses were inoculated to 72.1% (±5.6%) at 48 h when almost all *O. tauri* cells were lysed (Fig. 2 and Fig. 5).

**Figure 4.**
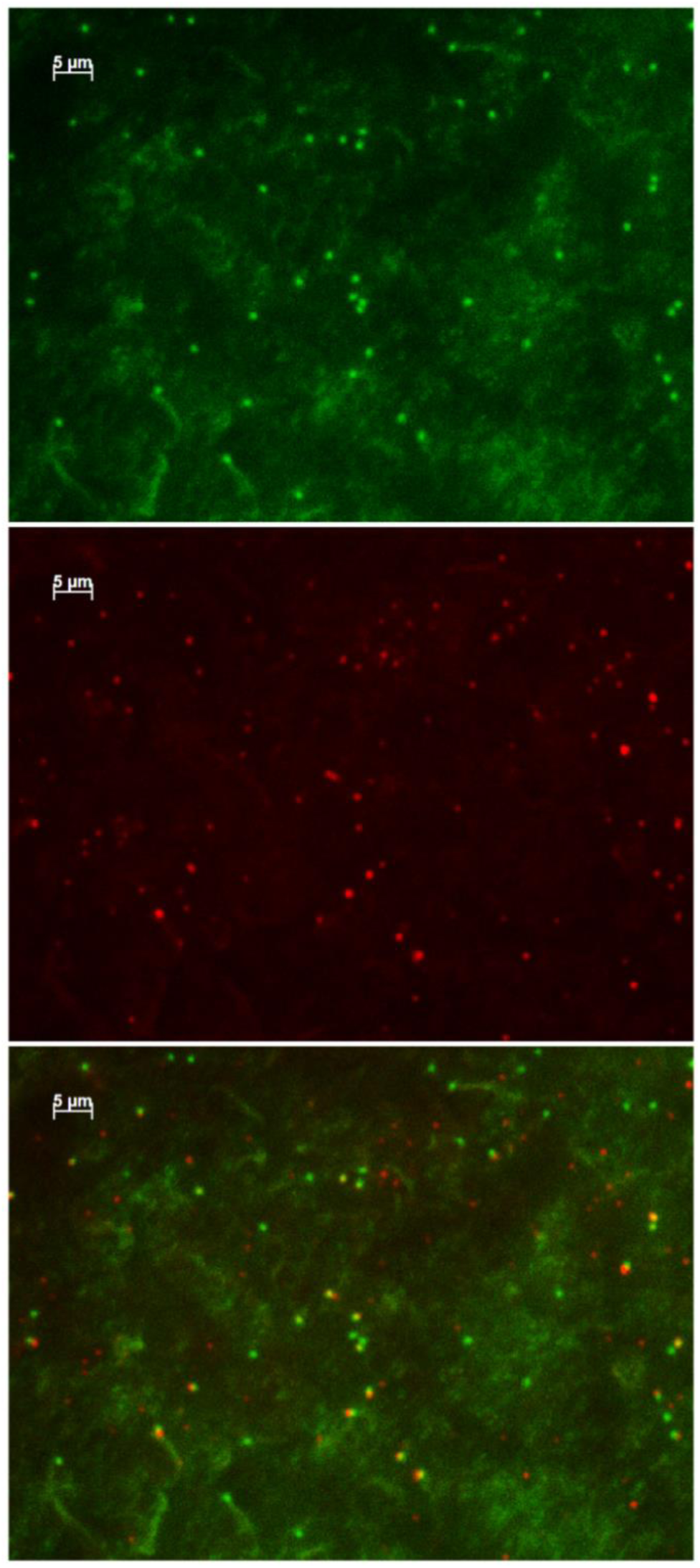
Micrographs of free viruses (at 48 h). Top: total viruses stained with SYBRGold. Center: VirusFISH labeled OtV5 viruses. Bottom: overlay of SYBRGold and VirusFISH signals for OtV5 viruses.

**Figure 5.**
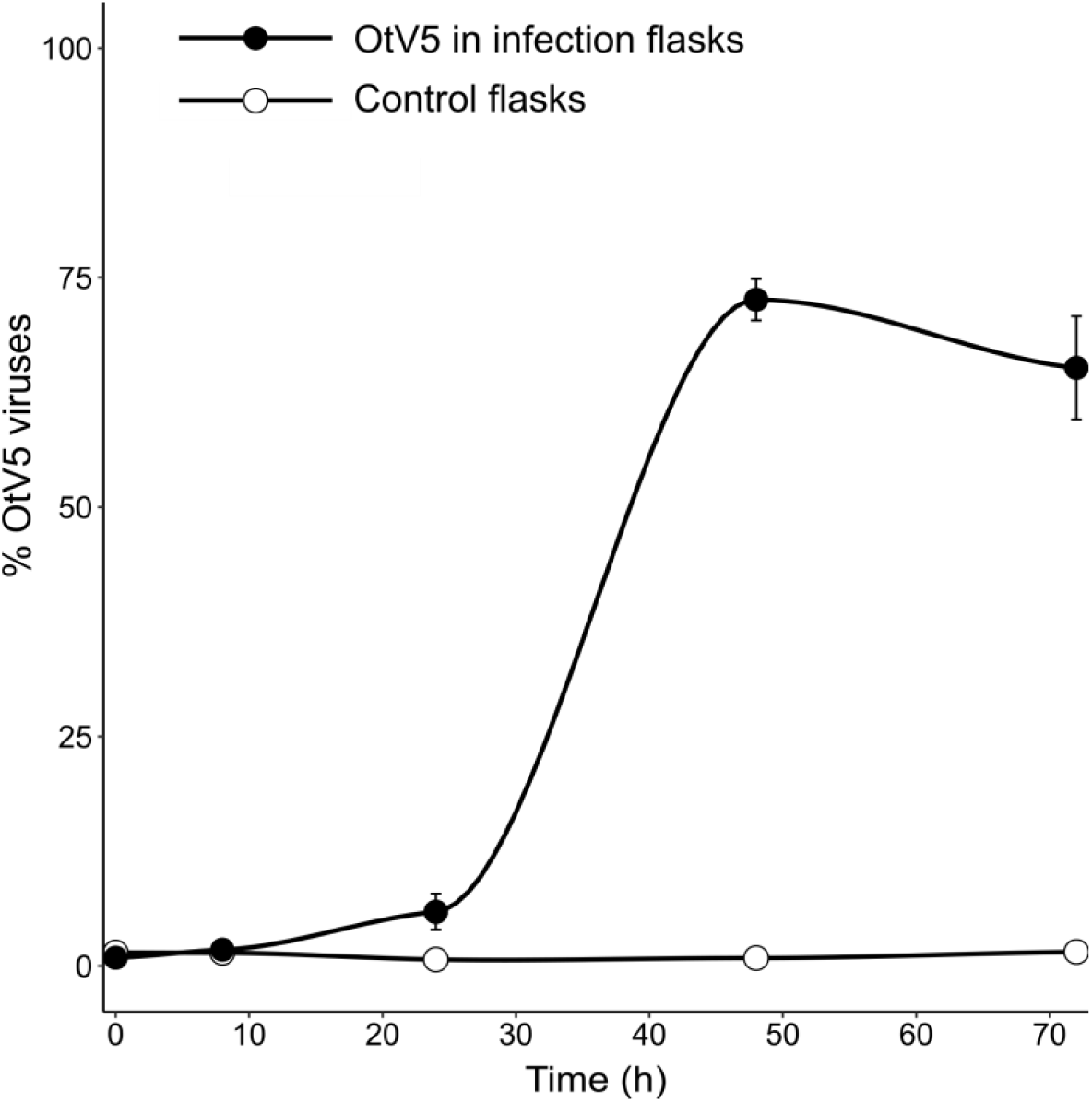
Dynamics of free viruses produced during the infection expressed as percentage of OtV5 with respect the total viral abundance. Counts were done by epifluorescence microscopy considering the overlay of signals.

### Burst size

Applying the formula developed by Middelboe & Lyck (2002) that considers the increasing and decreasing abundances of free viruses and their hosts, respectively, we obtained a burst size value of 7±0.4 viruses cell^-1^. To calculate this we considered that during lysis, the organic matrix released from the cells after 48 h trapped most of OtV5 particles on the filters (viral clouds). Therefore, to calculate the burst size we considered the number of free OtV5 particles at 24h (1.9·10^6^±3·10^4^ viruses mL^-1^) and, at 48 h, the sum of the number of free OtV5 (2.5·10^7^±3·10^6^ viruses mL^-1^) and the OtV5 particles within the viral clouds (2·10^8^ viruses mL^-1^) to obtain a final value of viral abundance at that time.

Moreover, we used VirusFISH to corroborate the burst size value obtained by the classical method, as mentioned in the Material and Methods section. For this, we used the observed average area of a single free OtV5 particle (0.13 µm^2^, n=30,000 viruses) from the Alexa594 fluorescence signal, and the average cellular area occupied by OtV5 virions at the maximum infection time-point before the major lysis occurred (24 h, 1.23 µm^2^, n=90 areas). Using this approach we obtained a value of 9.5±0.3 viruses per each infected *O. tauri* cell, which is very close to the results obtained for the burst size.

## DISCUSSION

Several studies have dealt with the virus—host relationships of the four clades of *Ostreococcus* spp. (*O. tauri, O. lucimarinus, O. mediterraneus* and clade B -Guillou et al., 2004-), and our knowledge on these systems is continuously expanding (Weynberg *et al.*, 2017). From these studies, only a few focused on the infection dynamics (e.g. Derelle *et al.*, 2008, 2017; Heath and Collins, 2016)(Derelle <i>et al.</i>, 2008, 2017; Heath and Collins, 2016), and most of the work has been directed towards understanding the virus–host interaction at the molecular level (e.g. Derelle *et al.*, 2008; Weynberg *et al.*, 2011; Clerissi *et al.*, 2012), unveiling interesting information on the host resistance mechanisms to viruses (Thomas, 2011; Heath and Collins, 2016; Yau *et al.*, 2016). However, to understand the impact of viruses on the ecology of *Ostreococcus* spp. it is crucial to develop techniques that enable monitoring the host—virus interactions at the single cell level, with the ultimate goal to apply them in complex natural communities. We designed probes to detect OtV5, but the alignment of the probes with other Prasinovirus genomes showed that they can very likely label all 11 genome sequenced *Ostreococcus* spp. viruses (Table S2 and Table S3), except OtV6, which is evolutionarily distinct (Monier *et al.*, 2017). Thus, our technique may help in fostering our knowledge on the role of viruses in the control of the abundance of the cosmopolitan *Ostreococcus* spp. Contrary to flow cytometry measurements and plaque-forming units assays, which only can give absolute cell and virus counts, VirusFISH allowed distinguishing and following the whole process of infection and shed light on what was happening previous to culture clearance, unveiling that infection was much more rapid than can be detected by cell or free virus counts. It showed that, despite most viruses seeming to have a period of latency after inoculation, some adsorbed producing a first discrete wave of infection after 8 h. At 24 h post-inoculation the infection percentage increased to a 16%, a quite low percentage if we consider that this process is followed by a surprising fast lysis of the culture only 24 hours later.

Another valuable application of VirusFISH was to determine the free viral particles released during infection, discriminating the true OtV5 from phages and other unspecific particles, improving the flow cytometry counts. Thus, we could estimate the burst size of an axenic culture (∼7 viruses per cell). Also, the technique allowed corroborating the obtained burst size results by estimating the amount of viruses inside the cell at late stages of infection, giving similar results (∼9.5 viruses per cell). If we compare these values with the 25 reported in Derelle *et al.* (2008) they do not extremely differ, despite there is a possibility that Derelle *et al.* (2008) could have overestimated the counts due to the fact that they used flow cytometry and could have counted phages as OtV5 particles. However, several studies have revealed that the experimental conditions affects the burst size value (Maat *et al.*, 2014; Maat and Brussaard, 2016), yielding a variation in the viral production among experiments. For instance, in *O. lucimarinus* (Zimmerman *et al.*, 2019) the infection of OlV7 virus differs depending on the growth light regimes. When the *O. lucimarinus* grows in optimal light conditions, the burst size is ∼680 virus/host, but when the light conditions are suboptimal, and thus the cellular machinery is not working properly, the burst size decreases to ∼50 virus/host. Therefore, burst size varies with the growing conditions. Nevertheless, burst size values may also vary depending on the physiological state of the cells, and may decrease in the stationary phase of the culture (Demory *et al.*, 2017).

Furthermore, although it was not the goal of our study due to the tiny size of *Ostreococcus*, VirusFISH could be potentially used for visualizing the dynamics of the viruses within the eclipse phase in larger hosts (i.e. nanoeukaryotes), something that is not feasible with other methods like Transmission Electron Microscopy.

### Methodological aspects to be considered for phototrophic eukaryotes and our particular O. tauri system

One of the best fluorochromes to label gene probes is Alexa594 (Barrero-Canosa *et al.*, 2017), which emits red fluorescence when excited with orange light. However, the chloroplasts of photosynthetic microbes also emit red fluorescence under the same light, hampering the detection of viral signals. We solved this technical issue by removing the cellular pigments with a combination of alcohol treatments, as described in the materials and methods section. The filter pore size also needs to be considered during VirusFISH experiments. *Ostreococcus* cells, although having a size of 1-3 µm, passed through a 0.6 µm filter, most likely because its cellular membranes are very flexible. This resulted in the loss of more than half of the cells during filtration. Consequently, we recommend the usage of filters with a pore size of 0.4 µm or 0.2 µm when working with picoeukaryotes. In our case, 0.2 µm pore size filters proved to be the best option, because, apart from completely retaining *O. tauri* cells, they allowed the visualization of viruses released from the lysed cells, and trapped in the organic matrix surrounding the cell debris (here referred as viral clouds) (Fig. 2, grey arrow). In contrast, these viral clouds could not be observed onto 0.4 µm filters, likely because the organic matrix passed through that pore.

### Modifications of VirusFISH with respect to the published protocols of phageFISH and direct-geneFISH

VirusFISH represents a combination between phageFISH and the direct-geneFISH. It used CARD-FISH to identify the unicellular eukaryotic host, similar to phageFISH, and used a mixture of polynucleotide probes directly labeled with a fluorochrome to target viral genes, similar to the direct-geneFISH protocol. CARD-FISH was used because its signal amplification step enables the detection of cells with low ribosome content. Indeed, *O. tauri* and all Mamiellophyceae have a small cytoplasm due to the big size of the organelles (Yau *et al.*, 2016), and therefore their ribosomal abundance is low and CARD-FISH enhances the cellular visualization. We also incorporated a step of embedding the filters in agarose to avoid cell losses in downstream manipulations of the filter portions. Furthermore, because *O. tauri* lacks a cell wall, the permeabilization step was omitted. On the other hand, a treatment to completely remove cell pigments was required, as mentioned above. Finally, compared to the direct-geneFISH protocol, we reduced the Alexa594 fluorochrome volume to label the viral gene probes in order to reduce economical costs but obtaining equal optimal results (see details in the methods section).

### VirusFISH vs other approaches to follow virus–host dynamics

Currently available methods to assess the dynamics between host and viruses during infection are i) the frequency of visibly infected cells (FVIC) (Wommack and Colwell, 2000), ii) Real Time PCR (RT-PCR) (Monier *et al.*, 2017) of viral genes and iii) the plaque assay, for counting plaque forming units (PFU) (Brussaard *et al.*, 2016). Compared with these methods, VirusFISH brings further advantages. For example, FVIC reports the fraction of infected host cells, but only detects those cells in the late stage of infection. PFU and RT-PCR describe the infection stages, but they lack the ability to measure the fraction of infected cells. With the exception of RT-PCR, which uses virus-specific primers, none of the three methods can identify the host or the viruses. In comparison, VirusFISH allows the: i) identification of both host and virus, using 18S rRNA and viral genes specific probes, feature particularly advantageous in non-axenic cultures of unicellular eukaryotes or in environmental samples; ii) quantification of the total and relative abundance of the host cells; iii) quantification of the total and relative abundance of virus infected cells, independent of the stage of infection and; iv) quantification of released viral particles. Furthermore, VirusFISH can potentially be used to discriminate the different stages of infection, in a manner similar to phageFISH.

Some other approaches have arisen in the last decade to unveil virus–host interactions, like the polony method (Baran *et al.*, 2018) or the microfluidic digital PCR (Tadmor *et al.*, 2011). The novel polony method is a culture independent technique based on a single molecule PCR. Using degenerate primers it allows the determination of the abundance of a given viral group and its degree of diversity, discriminating between different viral families or genera, and their host. This high-throughput approach has enabled the quantitative assessment of thousands of viruses in a single sample from both aquatic and terrestrial environments (Baran *et al.*, 2018). Thus, the polony method is a powerful approach to detect virus–host interactions in a cost-effective and relatively simple manner, but similar to VirusFISH, it requires the knowledge of the hosts and the genome of the viral target to design the probes. However, although the VirusFISH approach is not as high-throughput as the polony method, it has the advantage that it allows monitoring and visualizing a particular viral infection, so we can see when the infection is taking place, how many cells are infected at different times and how the infection progresses.

Also, VirusFISH allows quantification of the number of viruses that are being produced during an infection event, and the estimation of burst size values. Likewise, it allows distinguishing temperate virus infections (single viral copy detection in a cell) from lytic infections (when it detects multi-copies in a cell). Moreover, since VirusFISH consists on microscopy observations it enables the study of the heterogeneity of the infection within the host population, with the potential to extend its use to assessing those specific virus–host interactions in complex natural communities.

### Free viruses abundance and estimates of burst size values

The abundance of free viruses has been traditionally assessed through i) plaque-assay count (Suttle and Chen, 1992), ii) transmission electron microscopy (TEM) of uranyl acetate stained virus particles (Malenovska, 2013) and; iii) by epifluorescence microscopy (Hennes *et al.*, 1995) or flow cytometry (Marie *et al.*, 1999) of SYBRGreen stained viruses. Each of the above methods have limitations: i) the plaque-assay is constrained to cultivable hosts and viruses; ii) TEM is a time consuming and expensive technique and; iii) SYBR staining followed by epifluorescence microscopy or flow cytometry does not distinguish between infective and non-infective viruses, making it impossible to identify the virus of interest within a complex viral community. With VirusFISH we achieved the detection of specific free viruses in a relatively fast way, with no requirements of specialized equipment, or extremely expensive reagents. Allers et *al.*, (2013) also applied phageFISH to visualize free viral particles, by immobilizing the viral lysate on glass slides, which can potentially lead to virus losses. With VirusFISH we tried to overcome this issue by collecting and counting the free viruses on 0.02 µm anodisc filters decreasing the risk of viral losses during the hybridization process due to a better retention. The proportion of OtV5 in relation to all viruses present in the non-axenic culture, was 72% at 48h (Fig. 5), indicating that bacteriophages represented a minor fraction of the viruses at that time point. Later the proportion of OtV5 decreased slightly, probably due to an increase in the proportion of bacteriophages. This proportion would have likely been higher if we had taken samples after 72h, since organic matter released by the lysed cells fueled bacterial growth (data not shown).

In summary, in this study we developed VirusFISH to detect the virus–host interaction of a picoeukaryotic system. This technique allowed us to visualize and follow the dynamics of the OtV5 viral infection of *Ostreococcus tauri* until the complete lysis of the culture. Also, VirusFISH enabled the calculation of the viral production during infection, discriminating OtV5 viruses from the phages present in the culture. Moreover, we demonstrated that VirusFISH can be used to calculate the burst size of hosts in non-axenic cultures. Also, our designed probes could potentially target most *Ostreococcus* viruses, except for OtV6, representing a valuable tool to address virus–host interactions in these cosmopolitan marine picoeukaryotes. We strongly believe that VirusFISH presents great prospects to address infection dynamics in nature, and it will foster our understanding on the impact of viruses in eukaryotic populations. Furthermore, this technique can be easily implemented to any other model system.

## Supplemental Information

**Table S1.**
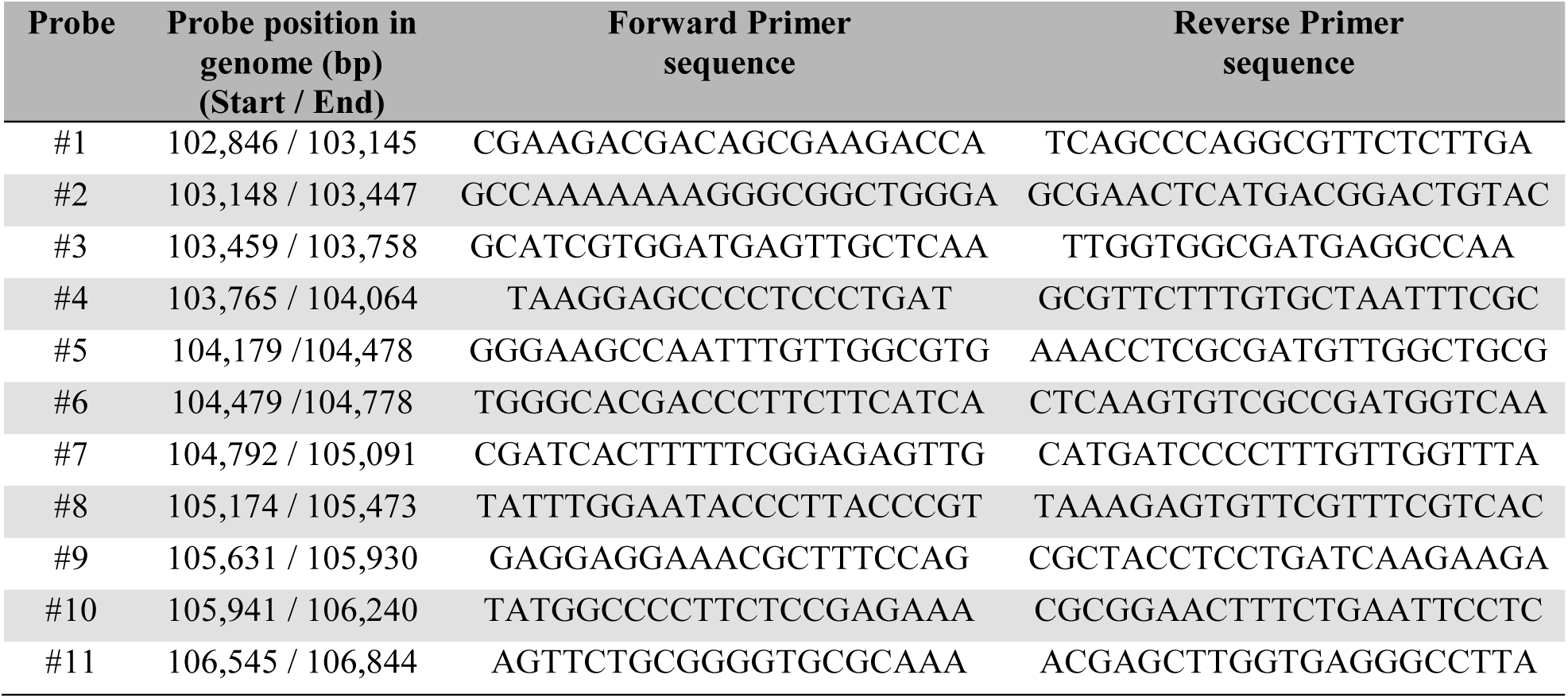
Position of the 11 probes in the OtV5 genome and the nucleotide sequence of the primers used to synthetize the viral probes.

**Table S2.** Alignment of our designed viral probes against Prasinovirus genomes and the outgroup PbCV1 (Chlorovirus) using BLAST. Shadow cells indicate identity higher than 80%, and coverage larger than 90%. Given than the probes are 300 bp long and all of them are combined for the hybridization, they may serve to visualize the virus-host interactions of all the *Ostreococcus* virus described in the table, except for OtV6. Accession numbers in Table S3.

|  | Probe 1 |  | Probe 2 |  | Probe 3 |  | Probe 4 |  | Probe 5 |  | Probe 6 |  | Probe 7 |  | Probe 8 |  | Probe 9 |  | Probe 10 |  | Probe 11 |  |
| --- | --- | --- | --- | --- | --- | --- | --- | --- | --- | --- | --- | --- | --- | --- | --- | --- | --- | --- | --- | --- | --- | --- |
|  | Id (%) | Cov (%) | Id (%) | Cov (%) | Id (%) | Cov (%) | Id (%) | Cov (%) | Id (%) | Cov (%) | Id (%) | Cov (%) | Id (%) | Cov (%) | Id (%) | Cov (%) | Id (%) | Cov (%) | Id (%) | Cov (%) | Id (%) | Cov (%) |
| OtV5 | 100 | 100 | 100 | 100 | 100 | 100 | 100 | 100 | 100 | 100 | 100 | 100 | 100 | 100 | 100 | 100 | 100 | 100 | 100 | 100 | 100 | 100 |
| OtV1 | 97.7 | 100 | 98.7 | 100 | 98.7 | 100 | 99 | 100 | 98.3 | 100 | 99.3 | 100 | 99.7 | 100 | 98.3 | 100 | 98.7 | 100 | 98.7 | 100 | 96.7 | 100 |
| OtV2 | 85.3 | 100 | 80.8 | 98 | 82 | 100 | 89.3 | 99 | 88.8 | 100 | 89.6 | 99 | 85.5 | 99 | 82.2 | 93 | 84.6 | 99 | 84.6 | 99 | 82 | 100 |
| OtV6 | 77.4 | 100 | 0 | 0 | 74.8 | 90 | 83.2 | 99 | 78.6 | 93 | 82.2 | 99 | 75.3 | 97 | 0 | 0 | 0 | 0 | 0 | 0 | 84.1 | 35 |
| OIV1 | 86.7 | 100 | 80.7 | 98 | 81.8 | 100 | 85.6 | 99 | 84 | 100 | 83 | 99 | 83 | 98 | 82 | 98 | 85.6 | 99 | 85.2 | 99 | 76.3 | 99 |
| OIV2 | 85.3 | 100 | 84.2 | 98 | 82.3 | 100 | 91 | 99 | 89.1 | 100 | 89.6 | 99 | 85.5 | 99 | 83.6 | 93 | 99 | 84.6 | 84 | 99 | 80.7 | 100 |
| OIV3 | 85.3 | 100 | 84.5 | 98 | 82 | 100 | 91.3 | 99 | 91.1 | 100 | 89 | 99 | 85.9 | 99 | 83.3 | 93 | 84 | 99 | 85 | 99 | 81 | 100 |
| OIV4 | 0 | 0 | 79.7 | 87 | 81.8 | 100 | 85.6 | 99 | 85.3 | 100 | 82 | 99 | 82.7 | 98 | 82.3 | 98 | 85.6 | 99 | 86 | 99 | 74.4 | 100 |
| OIV5 | 85 | 100 | 80.7 | 98 | 82.7 | 100 | 89.9 | 99 | 89.1 | 100 | 89.3 | 99 | 85.2 | 99 | 81.8 | 97 | 85 | 99 | 84.6 | 99 | 83 | 100 |
| OIV6 | 85.3 | 100 | 80.6 | 97 | 82.7 | 100 | 89.9 | 99 | 89.4 | 100 | 89.3 | 99 | 85.2 | 99 | 81.8 | 93 | 85 | 99 | 84.6 | 99 | 82 | 100 |
| OIV7 | 85.3 | 100 | 80.1 | 98 | 81.8 | 199 | 86 | 99 | 85.3 | 100 | 83.7 | 99 | 82.2 | 99 | 83.6 | 93 | 85 | 99 | 85.3 | 99 | 74.6 | 99 |
| OmV1 | 95.7 | 100 | 98.3 | 100 | 98.3 | 99 | 99.7 | 99 | 97.7 | 100 | 99.7 | 100 | 99 | 100 | 99.3 | 100 | 99.3 | 100 | 99.3 | 100 | 98.3 | 100 |
| OmV2 | 84 | 100 | 78 | 92 | 86.7 | 87 | 88.6 | 99 | 87.8 | 99 | 89 | 99 | 88.3 | 99 | 80.7 | 91 | 87 | 99 | 85.6 | 99 | 87.8 | 98 |
| MpV1 | 74.8 | 99 | 81.6 | 25 | 69.3 | 90 | 73.7 | 76 | 71.8 | 93 | 76 | 82 | 81.5 | 59 | 0 | 0 | 86.5 | 35 | 0 | 0 | 76 | 25 |
| BpV1 | 71.1 | 74 | 86.7 | 20 | 66.3 | 33 | 0 | 0 | 79.7 | 42 | 0 | 0 | 0 | 0 | 0 | 0 | 73.1 | 22 | 0 | 0 | 0 | 0 |
| PbCV | 0 | 0 | 0 | 0 | 0 | 0 | 0 | 0 | 0 | 0 | 0 | 0 | 0 | 0 | 0 | 0 | 0 | 0 | 0 | 0 | 0 | 0 |

Abbreviations: Id, Identity; Cov, Cover; OtV, *Ostreococcus tauri* virus; OIV, *Ostreococcus lucimarinus* virus; OmV, *Ostreococcus mediterraneus* virus; MpV, *Micromonas pusilla* virus; BpV, *Bathycoccus prasinos* virus; PbCV, *Paramecium bursaria Chlorella* virus 1.

**Table S3.** Genbank accession of Prasinovirus genomes and the outgroup PbCV1 (Chlorovirus) used in Table S2.

| Virus name | Accession number |
| --- | --- |
| OtV5 | EU304328.2 |
| OtV1 | FN386611.1 |
| OtV2 | FN600414.1 |
| OtV6 | JN225873.1 |
| OIV1 | MK514405.1 |
| OIV2 | KP874736.1 |
| OIV3 | HQ633060.1 |
| OIV4 | JF974316.1 |
| OIV5 | HQ632827.1 |
| OIV6 | HQ633059.1 |
| OIV7 | MK514406.1 |
| OmV1 | KP874735.1 |
| OmV2 | Described in a preprint article (Yau et al., 2019) and obtained from the authors |
| MpV1 | HM004429.1 |
| BpV1 | HM004432.1 |
| PbCV | NC_000852.1 |

Abbreviations: OtV, *Ostreococcus tauri* virus; OIV, *Ostreococcus lucimarinus* virus; OmV, *Ostreococcus mediterraneus* virus; MpV, *Micromonas pusilla* virus; BpV, *Bathycoccus prasinos* virus; PbCV, *Paramecium bursaria* *Chlorella* virus 1.

**Figure S1.**
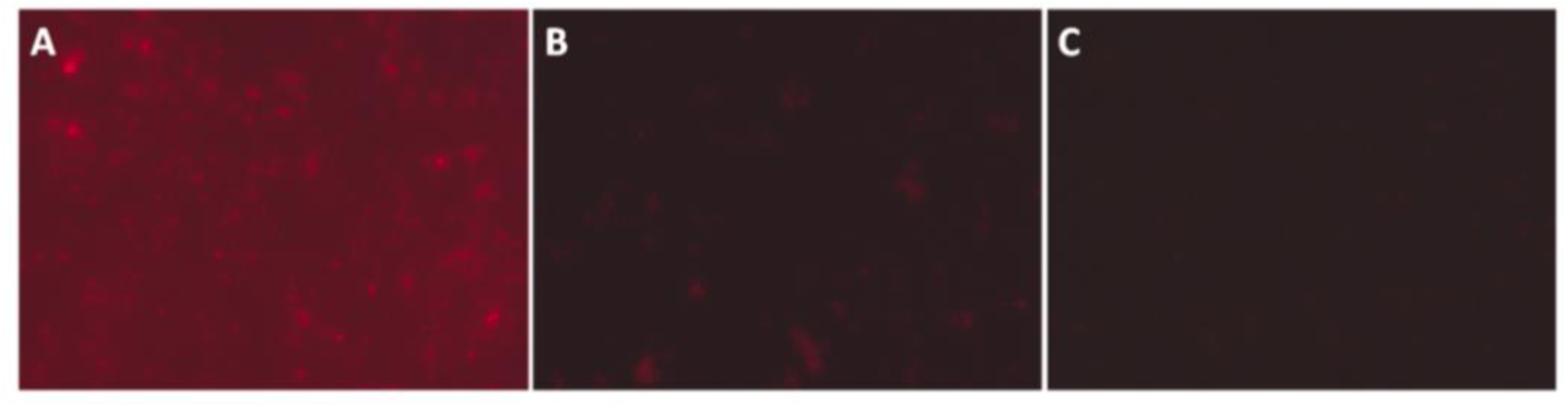
Cleaning test of *Ostreococcus tauri* 4221 culture for chlorophyll pigments removal. **A.** Negative control, no alcohol treatment. Chlorophyll is clearly visible. **B.** Cell culture treated for 1 hour with ethanol 97%. Chlorophyll is visually reduced, but not completely eliminated. **C.** Cell culture treated for 1 hour with ethanol 96% followed by 1 hour pure methanol. Chlorophyll is completely removed and almost no red background can be appreciated.

**Figure S2.**
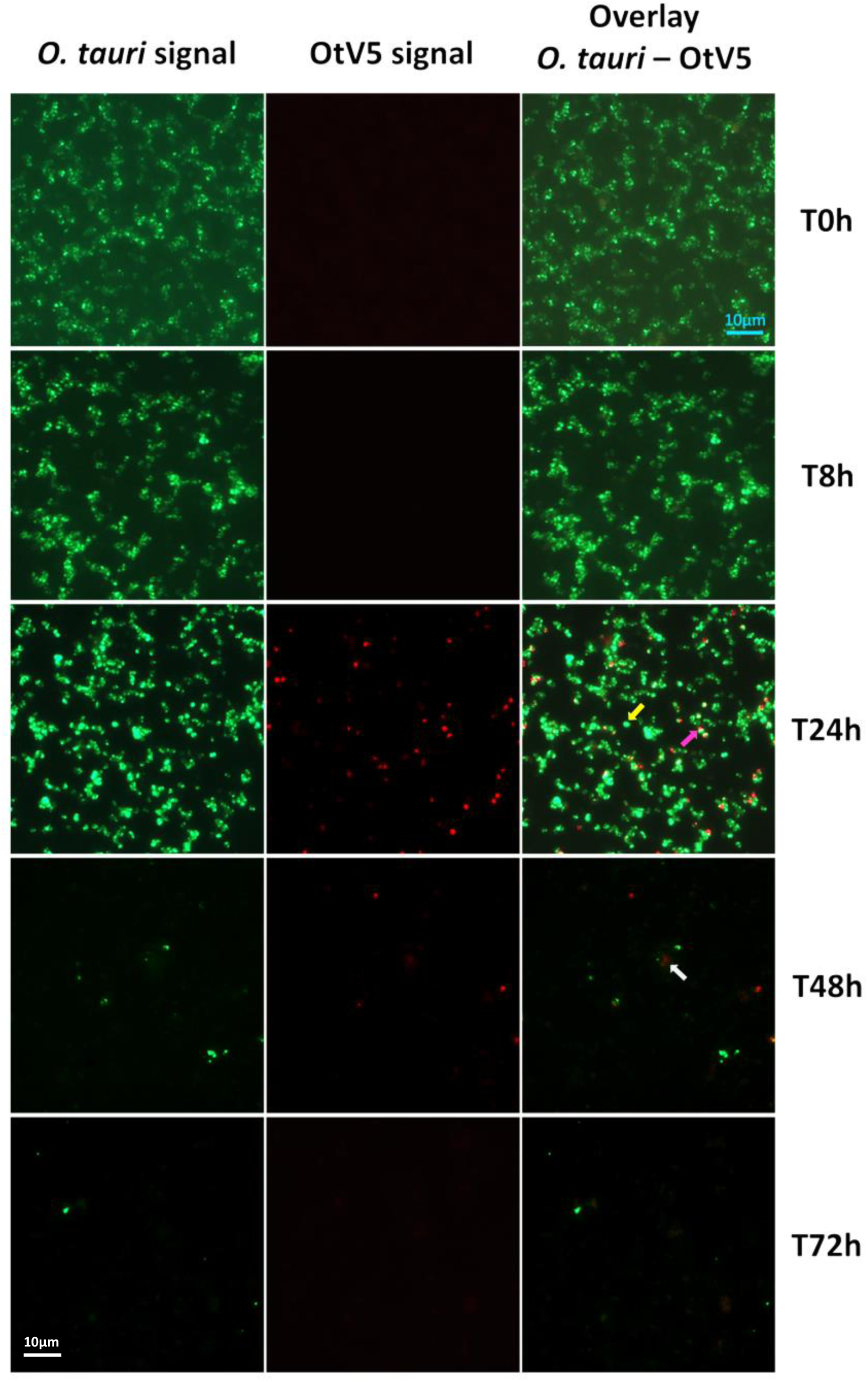
Micrographs of the evolution of the infection from time 0h to 72h. Left: *O. tauri* only. Centre: OtV5 only. Right column: overlay of *O. tauri* host cells in green (Alexa488) and virus in red (Alexa594). Yellow arrow: non-infected *O. tauri*; pink arrow: infected *O tauri*; grey arrow: cloud of viruses retained by the organic matter released during the lysis on the filter.

**Figure S3.**
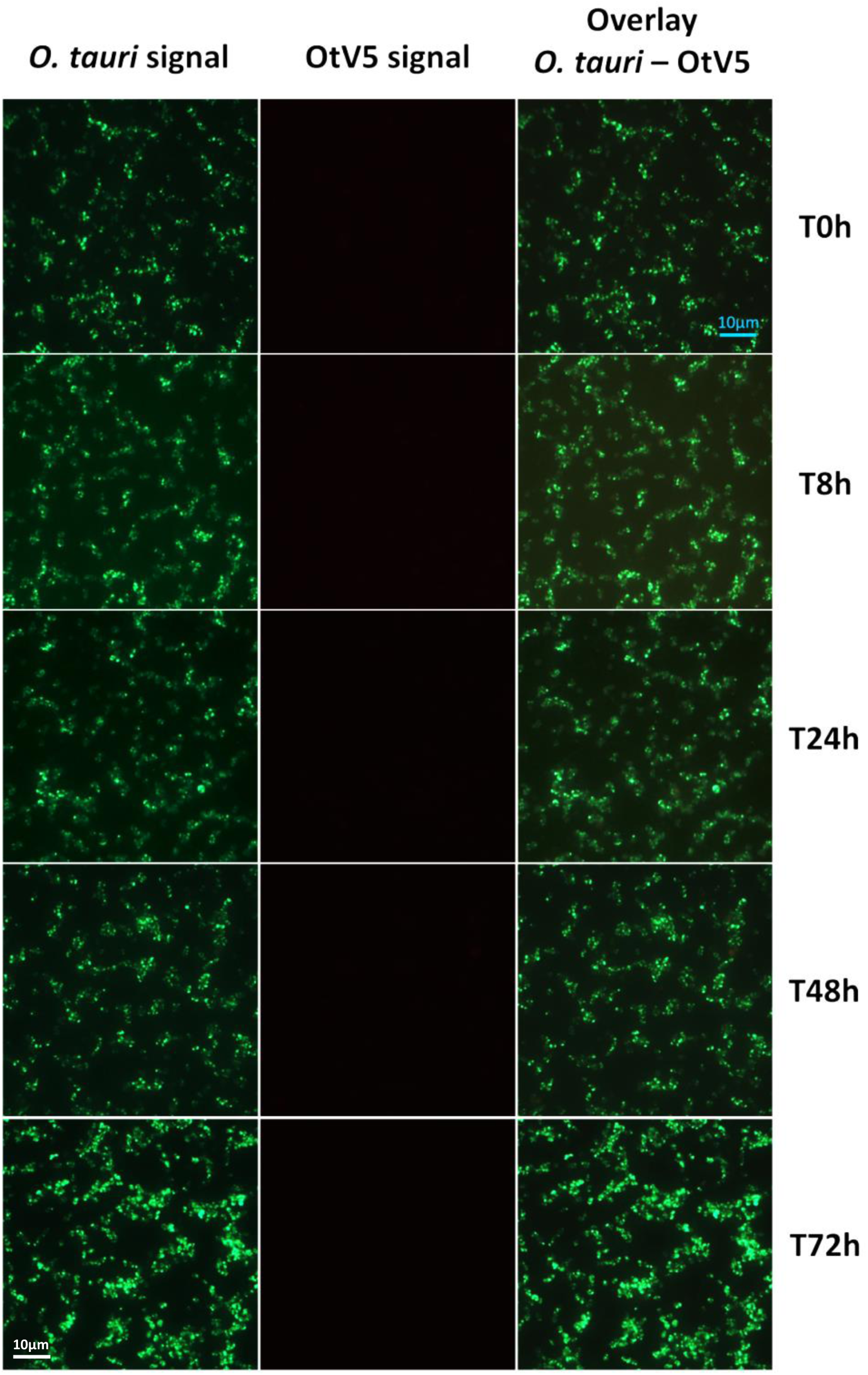
Micrographs of the evolution of the control flasks from time 0h to 72h. Left: CARD-FISH against *O. tauri* (Alexa488). Centre: VirusFISH against OtV5 viruses (Alexa594). Note no viral probes hybridation or false positives. Right column: overlay of both hybridization colors (Alexa488 and Alexa594).

**Figure S4.**
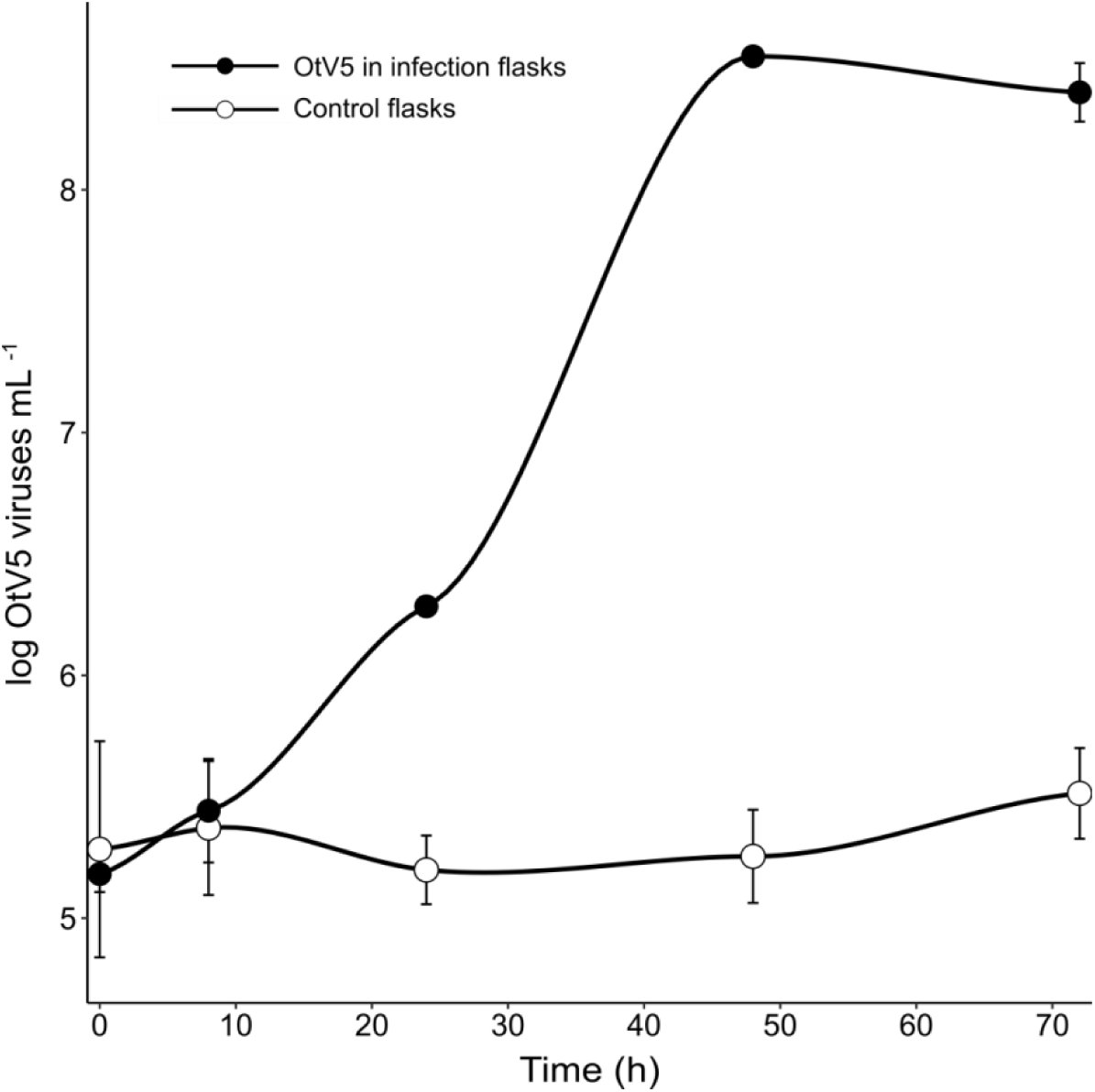
OtV5 particles abundance determined by VirusFISH on 0.02 µm anodisc filters. Black dots represent OtV5 particles in infection flasks. White dots represent the minimum basal false positive OtV5 viruses (mean of 3·10^5^ virus mL^-1^) detected in control flasks where no viruses were added.

